# Multimodal mucosal and systemic immune characterization of a novel non-human primate trachoma model highlights the critical role of local immunity during acute phase disease

**DOI:** 10.1101/2023.12.05.570239

**Authors:** Elodie Paulet, Vanessa Contreras, Mathilde Galhaut, Ida Rosenkrands, Martin Holland, Matthew Burton, Jes Dietrich, Anne-Sophie Gallouet, Nathalie Bosquet, Francis Relouzat, Sébastien Langlois, Frank Follmann, Roger Le Grand, Marc Labetoulle, Antoine Rousseau

## Abstract

**Background:** Trachoma -the leading cause of blindness worldwide as a result of infection-is caused by repeated *Chlamydia trachomatis* (Ct) conjunctival infections. Disease develops in two phases: i) active (acute trachoma, characterized by follicular conjunctivitis), then long-term ii) scarring (chronic trachoma, characterized by conjunctival fibrosis, corneal opacification and eyelid malposition). Scarring trachoma is driven by the number and the severity of reinfections. The immune system is a pivotal aspect of disease, involved in disease aggravation, but also key for exploitation in development of a trachoma vaccine. Therefore, we characterized clinical and local immune response kinetics in a non-human primate model of acute conjunctival Ct infection and disease.

**Methodology/Principal Findings:** The conjunctiva of non-human primate (NHP, Cynomolgus monkeys -*Macaca fascicularis-*) were inoculated with Ct (B/Tunis-864 strain, B serovar). Clinical ocular monitoring was performed using a standardized photographic grading system, and local immune responses were assessed using multi-parameter flow cytometry of conjunctival cells, tear fluid cytokines, immunoglobulins, and Ct quantification. Clinical findings were similar to those observed during acute trachoma in humans, with the development of typical follicular conjunctivitis from the 4^th^ week post-exposure to the 11^th^ week. Immunologic analysis revealed an early phase influx of T cells in the conjunctiva and elevated interleukins 4, 8, and 5, before a later phase monocytic influx accompanied by a decrease in other immune cells, and tear fluid cytokines returning to initial levels.

**Conclusion/Significance:** Our NHP model accurately reproduces acute trachoma clinical signs, allowing for the precise assessment of the local immune responses in infected eyes. A progressing immune response occurred for weeks after exposure to Ct, which subsided into persistence of innate immune responses. Understanding these local responses is the first step towards using the model to assess new vaccine and therapeutic strategies to prevent disease.

**Author Summary:** *Chlamydia trachomatis* is the leading infectious cause of blindness worldwide. The pathogenesis of trachoma is more complicated than other types of bacterial conjunctivitis: clinical signs of trachoma are rooted in repeated *Chlamydia trachomatis* infections of the inner eyelid surfaces, which roughens the skin. This lead to eyelid deformation and lashes rubbing on the cornea, which across multiple years of abrasion, ends with corneal opacification. The immune system is a pivotal aspect of disease, involved in disease aggravation, but also key for exploitation in development of a trachoma vaccine. Here we describe a non-human primate model of trachoma that accurately reproduces acute human eye disease, allowing for the precise assessment of the local immune responses in infected eyes. A progressing immune response occurred 4 weeks after exposure to Ct, which subsided into persistence of innate immune responses. Understanding these local responses is the first step towards using the model to assess new vaccine and therapeutic strategies to prevent disease.

## Introduction

Trachoma is currently the leading infectious cause of blindness worldwide. In 2022, trachoma was endemic in 42 countries (mainly in Africa), with 125 million people at risk, and 1.9 million visually-impaired [1,2]. The infection is caused by *Chlamydia trachomatis* (Ct), a Gram-negative obligate intracellular bacteria. The natural history of disease is divided into two successive phases: i) acute (or active) phase characterized by follicular conjunctivitis, and ii) scarring (or chronic phase) characterized by conjunctival fibrosis, eyelid malposition, and trichiasis ultimately causing corneal opacification [3,4]. Scarring or chronic trachoma develops over multiple years and after repeated Ct infections [5].

The *World Health Organization* (WHO)’s SAFE (*Surgery, Antibiotic, Facial cleanliness, and Environmental changes*) strategy, implemented in endemic zones by local ministries of health, efficiently reduced trachoma prevalence [6]. However, there are major drawbacks limiting long term efficacy of mass drug administration (MDA) preventive campaigns to combat trachoma. They include: i) off target antimicrobial resistance [7], ii) possible skewing of gut microbiome [8], iii) requirement of multiple rounds of treatment in some populations [5,9], iv) long term commitment of donor organizations to the control programs, v) a need for (and sometimes lack of) continued surveillance, and vi) absence of improvement in environmental and socio-economic conditions alongside control by MDA [10]. Altogether, a goal of eradication (rather than elimination) would only be achieved with a vaccine. Therefore, development of a vaccine strategy seems necessary to address these pitfalls in endemic areas [9,11]. Previous vaccine strategies have passed through different preclinical models and into human trials, but with mixed results [12–15]. A more precise understanding of trachoma pathogenesis and host-pathogen interactions are crucial for the identification of correlates of protection and the development of preventative strategies. While the extrinsic etiological agent of trachoma is Ct conjunctival infection, pathogenesis likely involves a substantial level of immune-mediated inflammation [3,4,16–18]. The crosstalk between Ct infection and the immune response during trachoma pathogenesis, is complex and undeciphered. However, in recent years significant advances have been made in understanding the immune response during trachoma pathogenesis [16]; with the role of non-specific and specific T cell responses being identified as essential for Ct clearance but concomitantly causing tissue damage [18]; neutrophil-mediated hyperinflammation has been shown in model infections to have negative consequences on ocular tissue damage and destruction [17,19]; and including changes in disease biomarkers, such as local cytokines during different trachoma stages [20]. However, studies are mostly cross-sectional, and published longitudinal data remains scarce, leaving open opportunities to improve comprehension of trachoma pathogenesis, which will be needed in the evaluation of a Ct vaccine. Improving our understanding of the precise mechanisms by which Ct triggers trachoma clinical and biological features requires longitudinal data in a relevant animal model. In this regard, NHP models are especially interesting due to their similarities with human immune responses and clinical manifestations [21,22].

We developed a NHP trachoma model that involved Ct ocular infection in macaques, which elicited follicular conjunctivitis that reproduced human features of acute trachoma [3,13,23]. We then characterized dynamics of local immune responses over time using a combination of multiparameter flow cytometry performed on conjunctival cells, tear and serum cytokine analysis, and Ct-specific immunoglobulin quantification.

## Methods

### Animals & Ethics statements

Six males (Group 1) and 12 females (Group 2) of Cynomolgus monkeys (*Macaca fascicularis*), aged 57 to 97 months for Group 1, and aged 36 to 38 months for Group 2, were used in this study. All animals were derived from Mauritius Island AAALAC-certified primate breeding facilities, and were housed in IDMIT’s animal facility (CEA, Fontenay-aux-roses, France, authorization #D92-032-02, Préfecture des Hauts de Seine, France) in individual cages during acute infectious phase, under BSL-2 containment.

General observations were conducted daily to assess well-being. Additionally, body weight and rectal temperature were monitored at each sampling time-point, along with a complete blood cell count (see Study Design). All experimental procedures were conducted according to European guidelines for animal care (“Journal Officiel de l’Union Européenne”, directive 2010/63/UE, September 22, 2010), and according to CEA institutional guidelines. The animals were anesthetized using intramuscular tiletamine (Zoletil 100®, 5mg/kg). Experimental protocols were approved by the institutional ethical committee (#17_072 (Group 1) and #21_008 (Group 2)), and were authorized by the “Research, Innovation, and Education Ministry” under registration number APAFIS#720-201505281237660 (Group 1), and APAFIS#31037-2021041418019968v1 (Group 2).

### Bacterial strain, exposure and treatment

B/Tunis-864 strain (B serovar) of *Chlamydia trachomatis* (Ct) was provided by the Statens Serum Institut (SSI, Copenhagen, Denmark). Bacteria were titrated and stored at -80°C at a concentration of 8x10^5^ infection forming units (IFU)/µL, then diluted in SPG buffer (sucrose 0.2M, sodium phosphate 20mM, and glutamic acid 5mM, pH 7.4) before administration at a final concentration of 2x10^4^ IFU/µL. Animals were inoculated with 20 µL of diluted bacteria using a micropipette, in both the inferior and superior conjunctival fornices of both eyes. Mock inoculated controls received only SPG buffer. At 7 weeks post-exposure (p.e.), animals in Group 2 received oral azithromycin (40 mg/kg, then 20 mg/kg daily, for 4 days).

### Study design

In a pilot study (Group 1), exposed animals to 10^4^ (Low dose, n = 2) or 10^5^ (High dose, n = 2) IFU/eye, or SPG buffer alone (SPG Buffer, n = 2), were followed with a standardized clinical follow-up throughout the study period, to define an appropriate challenge dose, and to determine a clinical grading scale. The exposure dose of 10^4^ IFU/eye was then selected for further characterization of immune responses, clinical symptoms, and bacterial load profiles in the Group 2 (n = 12) (**Fig 1**). All animals underwent identical standardized clinical follow-up throughout the study period.

**Fig 1.**
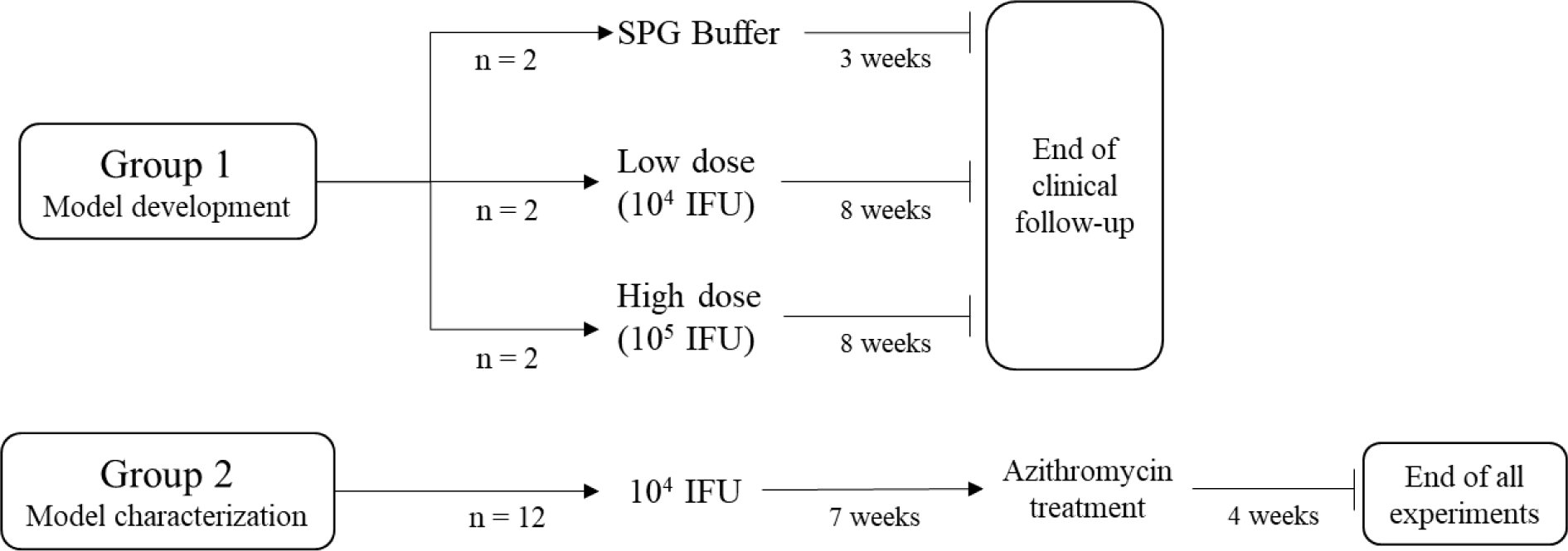
Study design. Group 1 was used to select an exposure dose while group 2 was used to characterize clinical manifestation, local and systemic immune response, and bacterial load.

### Conjunctival clinical scoring

Conjunctival clinical scoring was carried out using a customized clinical score assessed on ocular surface photographs (**Fig 2**). This score derived from the *WHO trachoma enhanced (FPC) grading scale* [24] and modified for compatibility with specific requirements of our model (**Fig 2**). Final score was calculated by adding the two components of the score: inflammation (graded on a scale of 0 to 3) and follicle (graded on a scale of 0 to 5), resulting in a maximum score of 8. Inflammation grading evaluates conjunctival edema, thickening, and the extent of hidden blood vessels. Follicle grading involves counting follicles in a predefined crescent-shaped zone of the upper tarsal conjunctiva (**Fig S1**).

**Fig 2.**
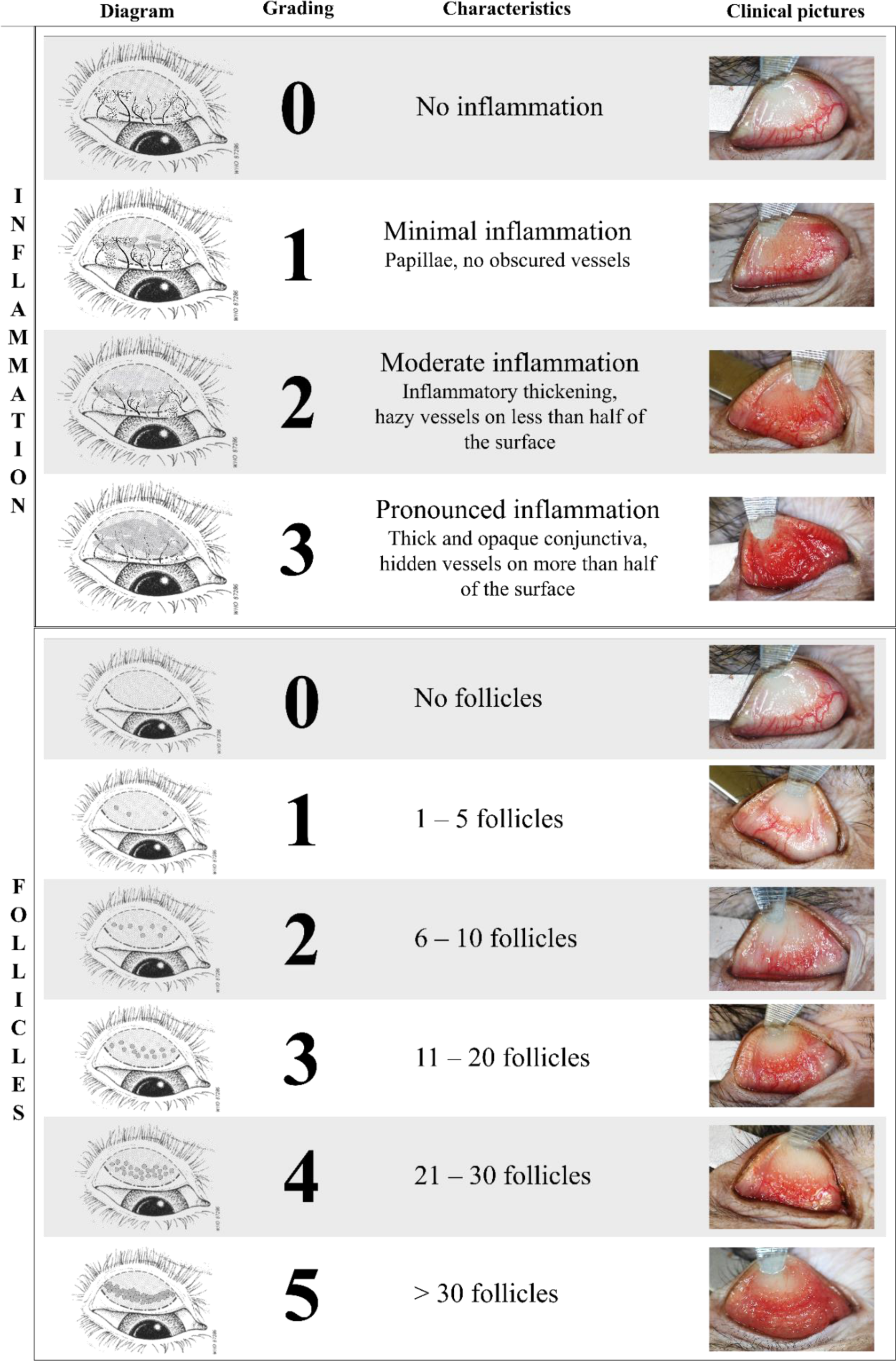
Conjunctival clinical scoring. Final score is calculated by adding the two components of the score: inflammation (graded on a scale of 0 to 3) and follicle (graded on a scale of 0 to 5), resulting in a maximum score of 8. Inflammation grading evaluates conjunctival edema, thickening, and the extent of hidden blood vessels. Follicle grading involves counting follicles in the central region of the upper tarsal conjunctiva. To obtain clear visibility, the upper eyelid was everted using a metal spatula and held in place with blunt forceps. Clinical photographs were captured while the animals were under general anesthesia. The schematic diagram of the upper tarsal conjunctiva was adapted from the World Health Organization© with permission.

Ocular surface photographs were captured following a standardized protocol, in brief: under general anesthesia, animal lying in supine position, upper eyelids were gently everted using a spatula, then held in place using blunt forceps (Dutscher, 956507).

An EOS 80D camera (Canon), equipped with a Macro 100mm Image stabilizer ultrasonic lens (Canon), and a Macro ring lite MR-14EXII circular flash (Canon), was mounted to a generic stand, facing down and positioned approximately 20cm above the eye of interest. Focus was set on the upper tarsal conjunctiva. Photographs were anonymized before scoring for inflammation grading and follicle counting by three independent observers masked to experimental group (A.R., E.P., M.G.), including one ophthalmologist trained and sub-specialized in human ocular surface infections (A.R.). Both eyes were scored independently by each observer. For each photograph, the scoring process involved the following criteria: i) if all three observers reached a consensus, the score was validated; ii) if two out of the three observers agreed, with the third observer differing by only 1 point on the clinical grading scale, the agreed-upon score was retained; and iii) if all three observers disagreed or if one observer had a score differing by more than 1 point with the two others, a second round of masked grading, involving all three observers, was conducted and the same criteria reapplied. As both eyes were inoculated with the same protocol, the average score of both eyes was used as the final score for each animal.

### Bacterial load quantification

Conjunctival swabbing was performed at the inferior conjunctival fornices instead of superior as to not influence clinical scoring and conjunctival imprinting. They were then placed in 1mL of room temperature Amies medium, from which DNA was extracted with QIAamp DNA Mini kit (Qiagen). Quantitative PCR was performed to assess the bacterial load using the Presto combined qualitative real-time CT/NG assay (Goffin Molecular Technologies, CG160100, and primer set previously described [25] CtPl+ 5’-TAGTAACTGCCACTTCATCA-3’ and CtP2 5’-TTCCCCTTGTAATTCGTTGC-3’) and using a CFX96TM real-time thermocycler (Bio-Rad)). The bacterial load, expressed in IFU-equivalent copies, was calculated by the thermocycler software, from a standard curve (obtained from 4x10^5^ to 4x10^0^ IFU/mL of bacterial stock strain B/Tunis-864 diluted in dPBS 1X). The limit of detection was fixed as 4 equivalent IFU/mL and limit of quantification at 40 equivalent IFU/mL.

### Cytokines quantification

Tears were collected with Dina strip Schirmer-Plus^Ⓡ^ tear test kits (Coveto, France), with the test strip placed on the inferior conjunctival fornices until no more impregnation is observed (generally 2-3 minutes). 50µL of NaCl were then added to the strip before being centrifuged (19,000 x g for 20 minutes, Sigma 3-16PK) to extract the tears. Bead-based Luminex^Ⓡ^ multiplex assay protocol (Milliplex Map, PRCYTOMAG-40K) was performed on 25µL of diluted tears (completed with PBS to obtain a total of 70µL) to quantify cytokine concentrations (G-CSF, GM-CSF, IFNγ, IL-1β, IL-1RA, IL-2, IL-4, IL-5, IL-6, IL-8, IL-10, IL-12/23(p40), IL-13, IL-15, IL-17α, MCP-1, MIP-1β, MIP-1α, sCD40L, TGF-α, TNF-α, VEGF, and IL-18). The same panel and protocol were applied to 25µL of undiluted serum samples for cytokine quantification.

### Conjunctival imprinting and flow cytometry

Local cellular immune infiltrates were evaluated through flow cytometric analysis of fluorescent antibody-labelled superficial conjunctival cell samples. Upper tarsal conjunctiva were gently dried with compresses. Imprints were harvested using 1.6 cm x 0.6 cm semi-oval pieces of nitrocellulose membrane (Supor® PES Membrane disc filters, 0.2 µm 47 mm, ref-66234, Pall, Ann Arbor, Michigan, USA), which were applied to the upper tarsal conjunctiva with gentle pressure. The membrane was then carefully removed and placed into phosphate buffer saline (PBS). After 1 hour of gentle agitation at RT, cells adsorbed onto the membrane were eluted using a flushing technique, as described previously [26]. Eluted cells were blocked with 10% rat serum before labelling with a panel of fluorescently-conjugated antibodies: CD326 (EpCam)-PE (clone 1B7, Thermo Fischer 12-9326-42), HLA-DR-AF700 (clone L243 G46-6, BD 560743), CD45-BV786 (clone D058-1283, Becton Dickinson 563861), CD66-FITC (clone TET2, Miltenyi 130-116-663), CD14-BUV737 (clone M5E2, Becton Dickinson 612764), CD3-BUV395 (clone SP34-2, Becton Dickinson 564117), CD8-BUV805 (clone SK1, Becton Dickinson 612890), CD4-BV421 (clone L200, Becton Dickinson 562842), CD20-PE-Cy7 (clone 2H7, Becton Dickinson 560735), CD159a(NKG2A)-APC (clone Z199, Beckman Coulter A60797). The samples were then fixed with BD Cellfix (Becton Dickinson 340181), and acquisition was performed using a 5-laser 27-filter ZE5 flow cytometer (Bio-Rad). Data were analyzed using FlowJo software (v10.8.1, Becton Dickinson). The gating strategy was designed to quantify immune subsets: neutrophils, monocytes, T lymphocytes, B lymphocytes, and natural killer (NK) cells (**Fig S2**). Leucocyte counts (CD45+ cells) were normalized for 50,000 observed cells, and all other immune populations were normalized for 100 leucocytes (to obtain proportions of immune cell populations expressed as percentage).

### Serum and tear IgG and IgA quantification

An indirect quantitative ELISA was used to measure specific anti-Ct IgG and IgA antibodies in serum and tears. Positive IgG and IgA references were arbitrarily assigned a value of 5 ELISA-Units/ml (AEU/ml). MaxiSorp plates (NUNC, Denmark) were coated overnight at 4°C with 4 μg/ml UV-inactivated B/Tunis-864 Ct elementary bodies in carbonate buffer pH 9.6. Plates were washed with PBS containing 0.05% Tween 20 and blocked with 2% bovine serum albumin (BSA) in PBS. Serum and tear samples were tested in duplicate, by 2-fold serial dilution in PBS with 1% BSA. IgA and IgG were detected by goat α-Human Fc IgA-Biotin conjugate and goat α-Human Fc IgG-Biotin conjugate (Southern Biotech, Birmingham, AL, USA) diluted 1:20,000 in PBS with 1% BSA and incubated for 1 hour at 37°C. This was followed by incubation for 40 minutes in a dark room with Poly-HRP40 (Fitzgerald, Acton, MA, USA) diluted 1:20,000 in PBS with 1% BSA. Detection substrate was TMB-PLUS (Kem-En-TEC, Denmark), the reaction was stopped with 0.5 M H_2_SO_4_ and absorbance was recorded at 450 nm (after subtraction of the background absorbance value measured at 620 nm). The IgG and IgA references were then used to establish a standard curve for the determination of titers in arbitrary ELISA Units/ml (AEU/ml) based on a five-parameter logistic curve using the package ‘drc’ in R (version 4.3.1) [27].

For determination of total IgG and IgA in tears, the same ELISA protocol (MaxiSorp, Nunc) was used except that plates were coated with anti-human kappa and lambda light chain specific mouse antibodies (Southern Biotech) at 1:1 ratio diluted to 1 μg/ml in PBS overnight. Purified human IgG and IgA were used as standards (Sigma, St. Louis, MO, USA). Tear samples were tested by 4-fold serial dilution, and serial 2-fold dilutions of IgG and IgA standards were applied. Total IgG and IgA were calculated from the IgG and IgA standard curves based on a five-parameter logistic standard curve.

### Statistical analysis

Correlation between multiple parameters was assessed using the non-parametric Spearman correlation test, with a two-tailed p-value, performed using R software (version 4.3.1) [27]. For each combination of two parameters, this analysis assigned a correlation factor (r) from -1, a maximum negative correlation to 1, a maximum positive correlation, and the p-value. For all other statistical analyses, particularly to confirm significant differences during longitudinal observation, the Wilcoxon test as well as the Friedman test (with a two-stage linear procedure correction) were conducted using Prism (GraphPad version 9.5.1).

## Results

### Clinical manifestations of the conjunctival infection

Two doses of inoculum were initially tested in a pilot study (**Fig 1**) to determine the lowest dose of Ct that gave disease in 100% of animals. From week 1 post infection, both animals in both groups exhibited clinically-defined acute trachoma, characterized by the presence of combined conjunctival follicles and inflammation (**Fig S3**). SPG Buffer mock-inoculated controls maintained a conjunctival clinical score ≤ 2 for the follow-up period (**Fig S3**). Subsequent experiments were conducted using the lower dose inoculum (10^4^ IFU) as it could elicit measurable disease in both animals.

Clinical conjunctival scoring and bacterial load quantification were performed for up to 11 weeks in monkeys (n = 12, MF1-12) infected with 10^4^ IFU of Ct (Group 2). At baseline, follicles (11/12, **Fig S4A**) and inflammation (10/12, **Fig S4B**) were almost completely absent. The outliers for follicles or inflammation were mild (scoring 1 and 0.5, for inflammation by MF5 and MF6, respectively, and a 0.5 follicle score for MF2). Follicular conjunctivitis was observed in all animals from 2 weeks post-exposure (p.e.) (12/12, p<0.0001), with 8/12 also demonstrating inflammation (median score = 2) (**Fig S4A and B**). Clinical scores peaked at 4 weeks p.e. and then steadily decreased (**Fig 3A**). Although at 6 weeks p.e., higher than the average clinical scores were recorded for almost half of the group (n=5 animals) until the end of the study. At week 6, inflammation score decreased for 5/12 animals. However, median clinical scoring decreased *post* azithromycin treatment (p.t.), as 10/12 animals demonstrated decreases in inflammation (**Fig S4B**).

**Fig 3.**
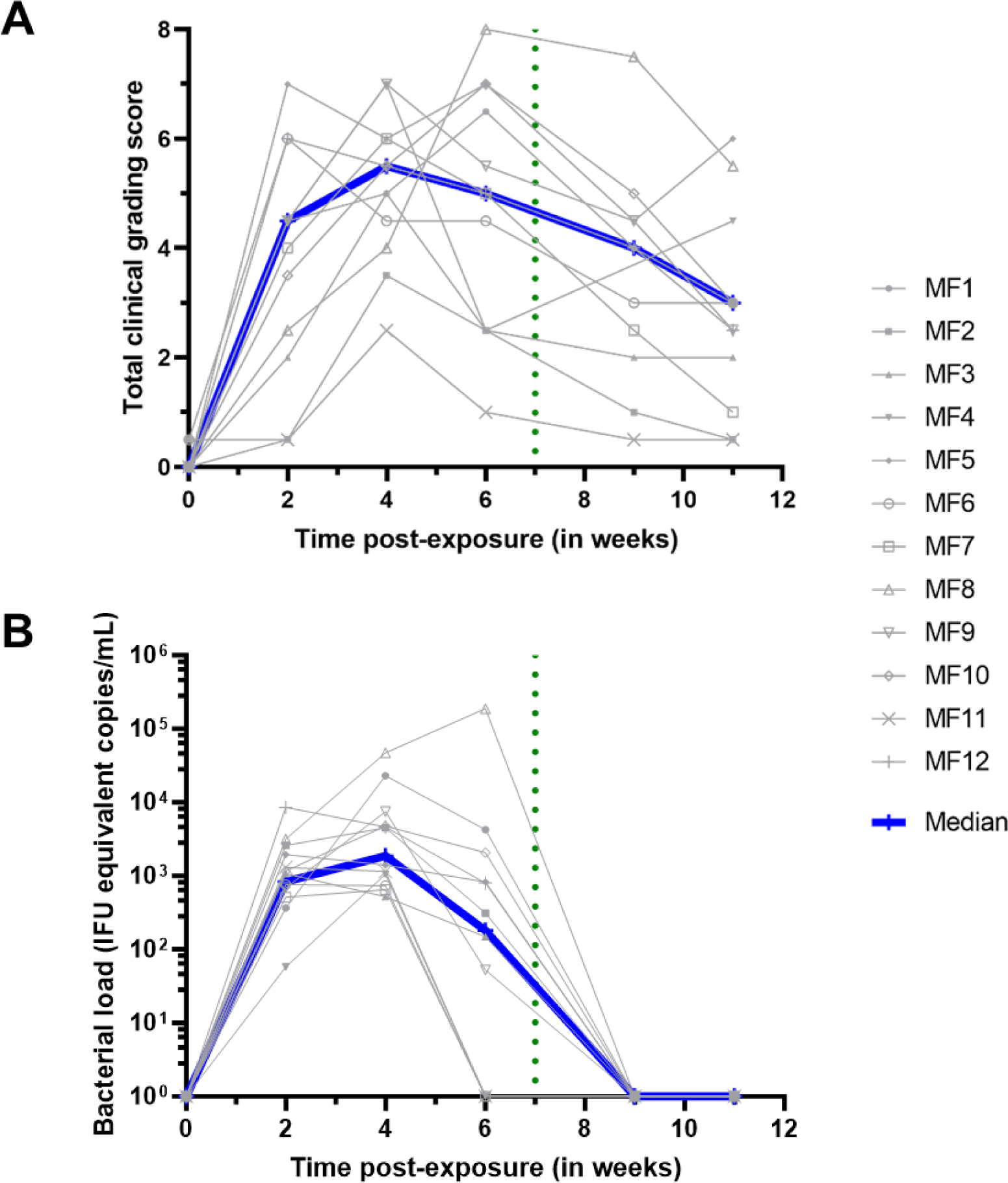
Ocular manifestations and bacterial load follow-up. The baseline was adjusted to week 0. All animals were infected with *C. trachomatis* (Ct) at week 0. The green dotted line represents the azithromycin treatment. (A) Ocular clinical grading scores were assessed using the *Clinical Grading Scale* (**Fig 2**) by three masked observers. The scores represent the average of both eyes and the three observers. The total clinical grading score for group 2 was determined by combining the inflammation and follicular scores (individual scores can be found in **Fig S4**). The Friedman test, with a two-stage linear step-up procedure correction by Benjamini, Krieger, and Yekuteli, confirmed significant changes over the course of the study. (B) Bacterial load quantification is expressed in IFU equivalent copies/mL. Experiments were performed for group 2 by conducting qPCR on DNA extracted from ocular fluids.

All other clinical parameters, including weight, temperature, and complete blood count, remained stable throughout the study, except for blood neutrophils, which increased following exposure to Ct (**Fig S5**).

### A robust conjunctival Ct infection

Conjunctival infection was monitored by changes in bacterial load, characterized by Ct-specific genomic qPCR (**Fig 3B**). All animals harbored Ct infection, which was detected at the first time-point of testing (2 weeks p.e.). Peak bacterial load was measured mainly 4 weeks p.e., and decreased in all animals except in one (MF8), which kept rising until after p.t.. In 4/12 animals, the bacterial load dropped to undetectable levels at 6 weeks p.e. In the 8/12 remaining animals with detectable load at week 6, bacterial load in all eyes dropped to undetectable levels by week 9, which was 2 weeks p.t. (**Fig 3B**).

These data demonstrate robust conjunctival Ct infection of all monkeys to produce scorable clinical disease.

### Assessment of conjunctival immune responses

### Immune cell populations at the ocular surface

Flow cytometry was performed on conjunctival cells sampled during the course of infection, to assess the major constituents of the local immune response. Leucocyte numbers (CD45^+^ cells normalized to 50,000 observed cells) had increased at 1 week p.e., and peaked mainly at 3 weeks p.e. (*p = 0.0078* baseline vs week 3 p.e. (**Fig 4A**). The most common leucocytes detected by sampling were T cells, monocytes, and “other leucocytes”, defined as CD45^+^ cells negative for all other immune markers (**Fig S2**). During the first 2 weeks p.e., the proportion of T cells increased slightly from baseline (57 ± 8% to 67 ± 9%, *p = 0.1210*), and remained stable (**Fig 4B**). Between weeks 3 and 4, T cells proportions fluctuated and by week 6 this compartment had contracted. Interestingly, between weeks 3 and 6 p.e., B cells (CD20^+^ HLADR^+^-double positive CD45^+^ cells) were detected in significant proportions, with the highest proportion at 6 weeks p.e. (2.14 ± 0.57%, p = 0.0156 week 6 vs week 1 p.e.) (**Fig 4B**). The frequencies of monocytes fluctuated from week 1 to week 4 p.e. without significant differences from baseline (*p = 0.8750*), then their proportion among leucocytes steadily increased from 6 weeks p.e., with the highest proportion observed at 11 weeks p.e. (53 ± 9% of all immune cells, *p = 0.0078* week 11 (final time tested) vs week 1 p.e.). Negligible proportions of neutrophils (CD66^+^ cells) and natural killer (NK, CD159a-NKG2A^+^ cells) cells were observed throughout the study period (less than 1% of immune cells).

**Fig 4.**
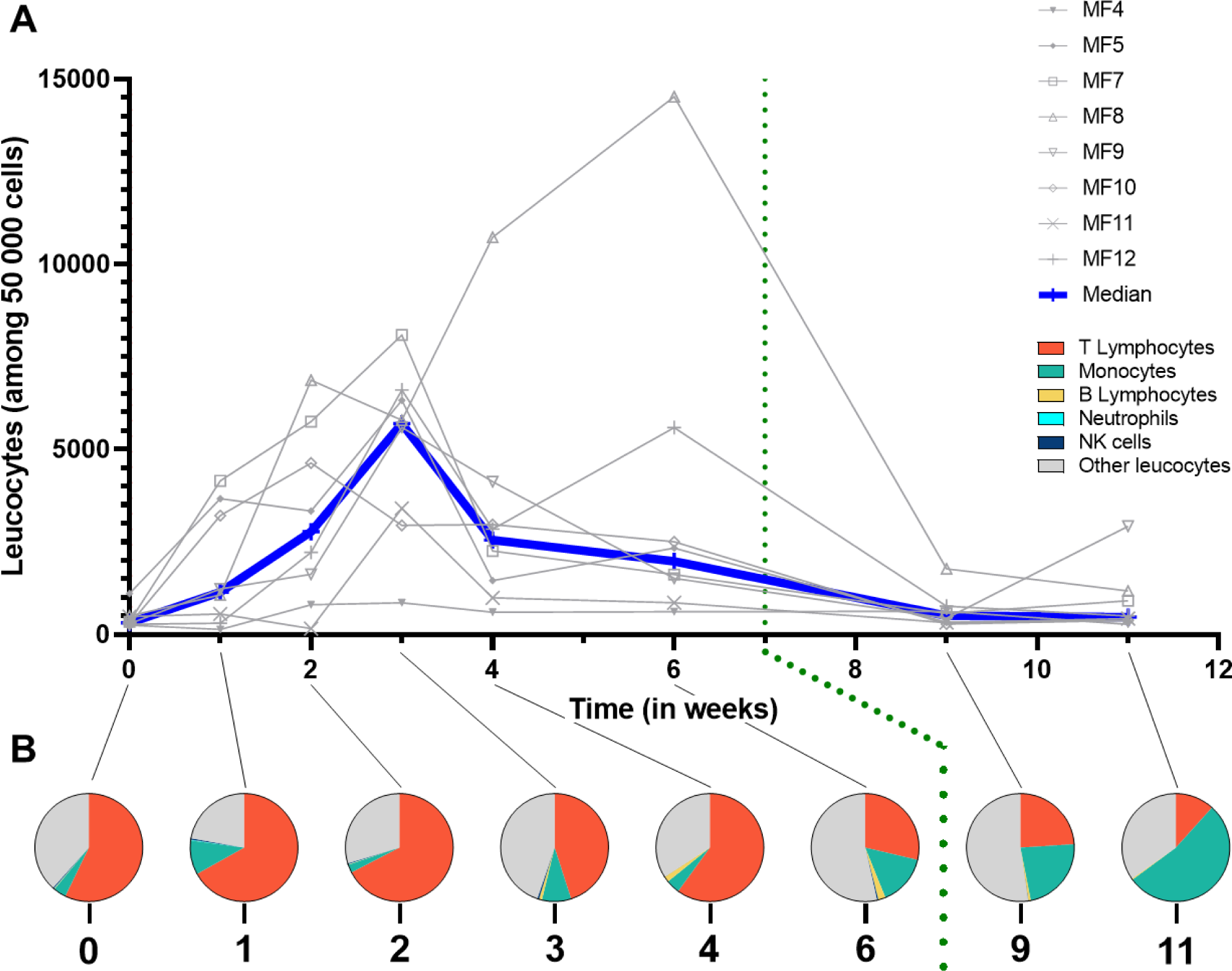
Multiparametric flow cytometry of superficial conjunctival cells. (A) Leucocytes count normalized for 50,000 cells. (B) Specific immune population proportion normalized for 100 leucocytes. The baseline was adjusted to week 0. All animals were infected with *C. trachomatis* at week 0. The green dotted line represents azithromycin treatment. Other leucocytes category represents non-identified immune cells. The Friedman test, with a two-stage linear step-up procedure correction by Benjamini, Krieger, and Yekuteli, confirmed significant changes over the course of the study.

These data demonstrate local fluctuations of key immune subsets responding to acute Ct infection and its treatment.

### Dynamics of cytokine secretion in response to Ct at the conjunctiva

To explore local immune responses further, ocular surface cytokines were quantified using Luminex (**Fig 5A**). Most cytokines were consistently detected from the conjunctiva of animals before infection, at baseline. Some cytokines (TNF-α, MIP-1α, IL-6, IL-18, and G-CSF) concentration did not significantly change after exposure to Ct (**Fig S6**). Cytokines VEGF, IL-12/23, IL-13, IL-17α, IL-10, IFN-y, IL-2, IL-15, sCD40Land IL-1β decreased between weeks 4 or 6, or during both, but rebounded at week 11 p.e., to higher levels than original baselines, and in more animals. Meanwhile, the pattern of detection was different for TGF-α, MIP-1β, IL-8, MCP-1, and IL-1RA, which increased at some point between weeks 2 to 6 p.e. (p=0.0239, 0.0003, 0.0068, 0.0239, and 0.0049 at peak respectively), but then subsided by week 11. Finally, a fourth pattern was observed with cytokines IL-4, IL-5 and GM-CSF remaining low through the course of the experimental period, then increasing drastically only at week 11. These results demonstrated 4 different conjunctival cytokine response patterns to Ct, which either did not change significantly, increased or decreased in correlation to bacterial load, or increased following treatment when bacterial load was abolished.

**Fig 5.**
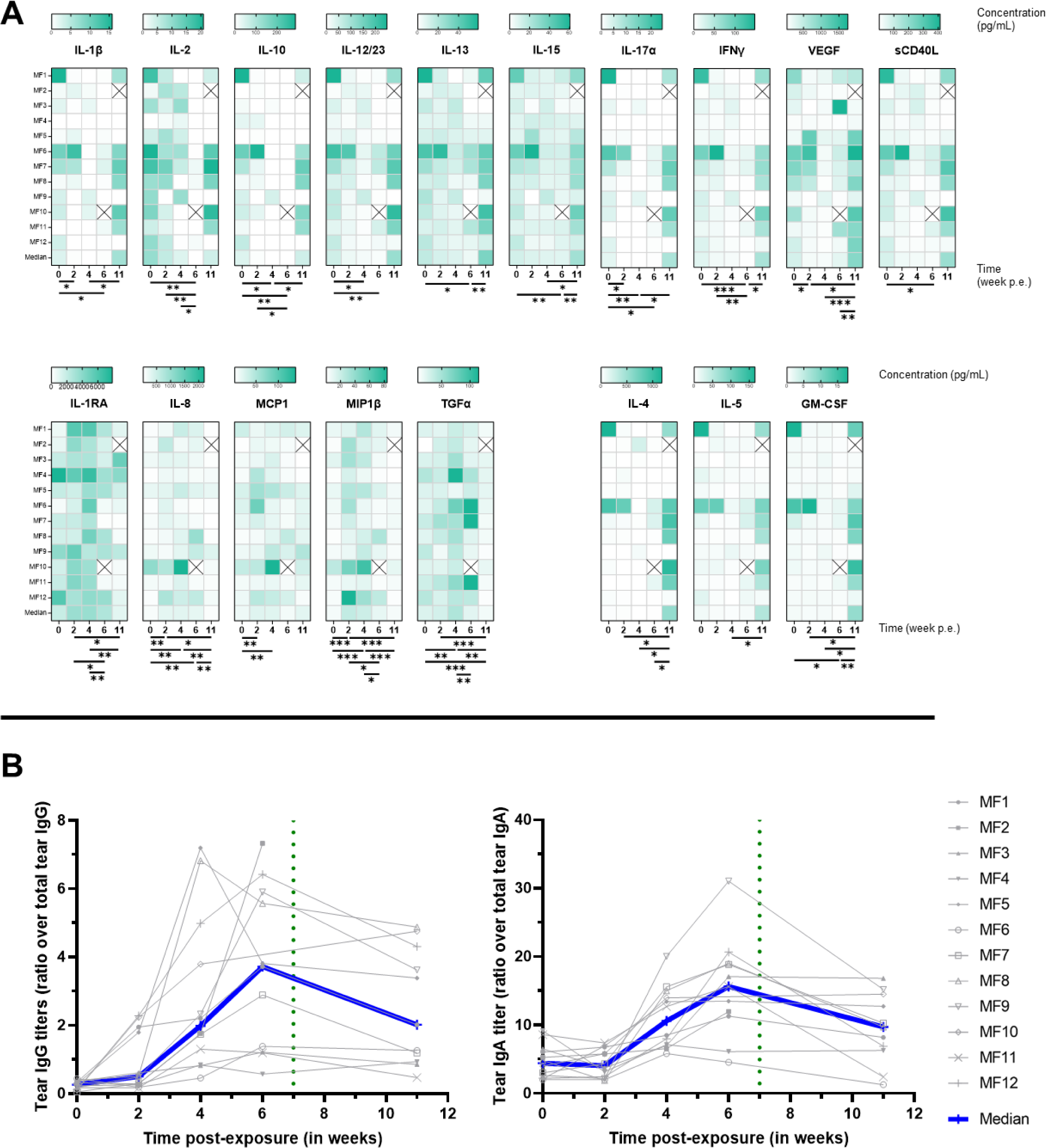
Cytokines and Ig secretion in tears. The baseline was adjusted to week 0. (A) Cytokines quantification performed by Luminex on tears. The Friedman test, with a two-stage linear step-up procedure correction by Benjamini, Krieger, and Yekuteli, was utilized to confirm significant changes over the course of the study. **** p<0.0001, *** 0.0001<p<0.001, ** 0.001<p<0.01, * 0.01<p<0.1. (B) IgG and IgA quantification performed by specific ELISA assay on tears. Results are shown as a ratio of Ct-specific tear Ig over total tear Ig.

### Increased local IgA and IgG in response to Ct exposure at the conjunctiva

IgA and IgG specific to Ct elementary bodies were quantified from tears to determine the local adaptive humoral response to conjunctival Ct exposure. IgG increased comparably at 2 weeks p.e. (p=0.021), then both IgG and IgA peaked between 4 and 6 weeks p.e. (p=0.001 and 0.002 between weeks 0 and 6 for IgG and IgA respectively), and then decreased at 11 weeks p.e. (p=0.0098 and 0.0039 for IgG and IgA respectively) (**Fig 5B**). These data demonstrated that local adaptive humoral responses developed over the first 6 weeks p.e.

### Systemic response to conjunctival Ct exposure

Serum cytokines and specific IgG and IgA were measured also from sampled serum. Interestingly, the pattern of TGF-α detection in serum matched the fluctuations observed in tears, increasing to peak at 4 weeks p.e., then subsiding by week 11 (**Fig 6A**). In contrast, sCD40L continued to increase in serum at each time-point tested, to peak at week 11 p.e.. IL-8 in serum decreased by 4 weeks p.e., only to rebound to higher levels than initial baseline levels at week 11 (the inverse to detection pattern in tears). MIP-1β detection pattern in serum mimicked that of tears, increasing by week 4 and subsiding by week 11 (**Fig 6A**). Those results suggest that, while limited, sCD40L, TGF-α, and MIP-1β secreted by different immune cells may play a role in early systemic response to Ct exposure, while IL-8 could be implicated in the later phase of infection

**Fig 6.**
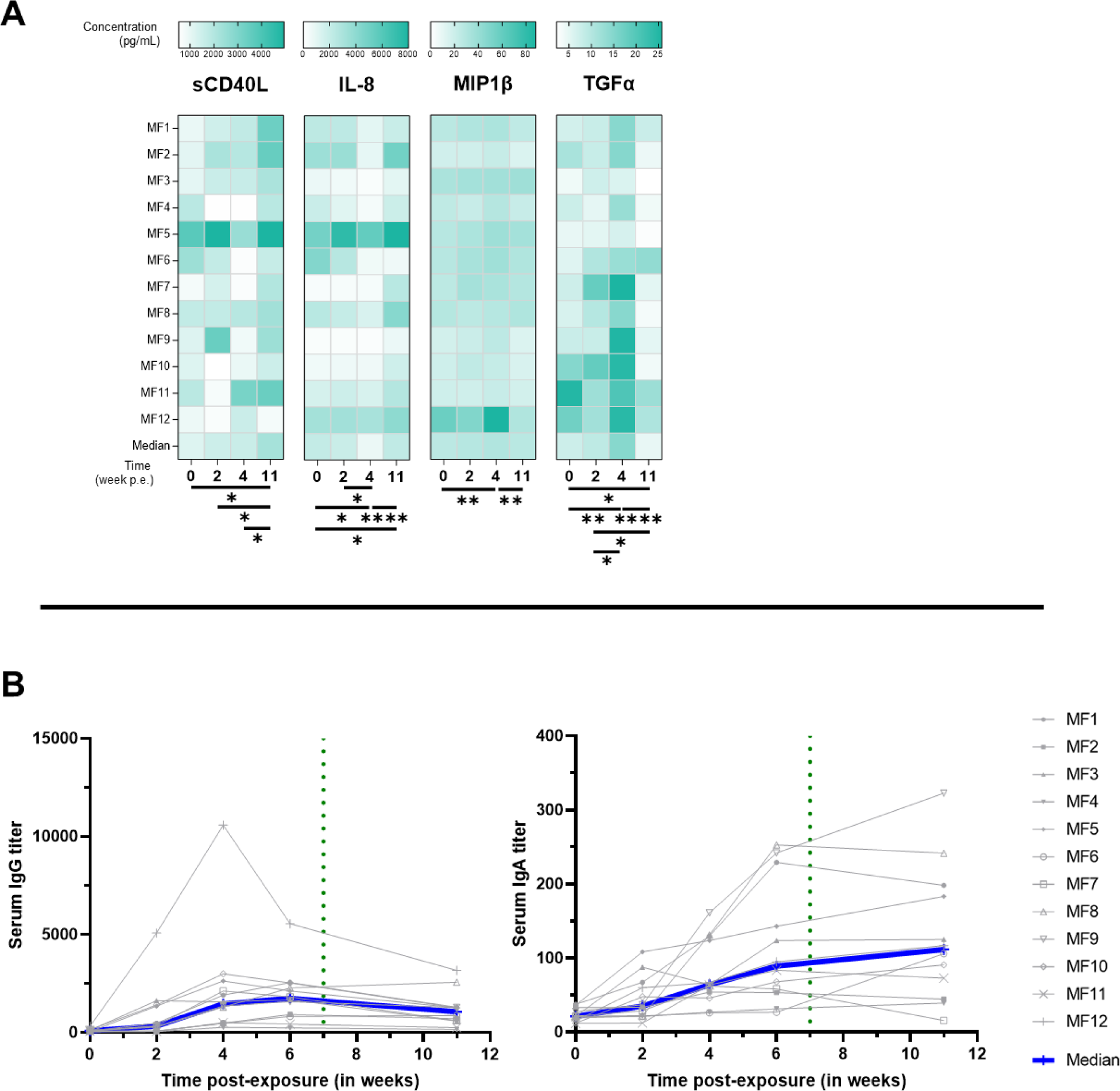
Cytokines and Ig secretion in serum. The baseline was adjusted to week 0. (A) Cytokines quantification performed by Luminex on serum. The Friedman test, with a two-stage linear step-up procedure correction by Benjamini, Krieger, and Yekuteli was utilized to confirm significant changes over the course of the study. **** p<0.0001, *** 0.0001<p<0.001, ** 0.001<p<0.01, * 0.01<p<0.1. (B) IgG and IgA quantification performed by specific ELISA assay on serum.

Additionally, circulating Ct-specific IgA and IgG quantification was performed on serum samples (**Fig 6B**). Increases in IgG were detected in 7/12 animals p.e., peaking between weeks 4 and 6 and declining subtly by week 11. Ct-specific IgA increased in 10/12 animals from 2 weeks p.e., but unlike IgG, continued to rise between week 6 to week 11. These results demonstrate that Ct exposure at the conjunctiva elicits systemic circulation of Ct-specific IgA and IgG, which remain elevated at the latest time-point tested.

### Biomarker signature associated with clinical scoring and bacterial load

All of the parameters analyzed above, at all time-points, were assessed for correlations using the Spearman correlation test (**Fig 7**). This combined analysis uncovered 2 distinct signatures: a positive or a negative correlation with bacterial load. Increases in clinical signs (inflammation, follicles and total clinical signs) correlated with each other, as well as with bacterial load (p<0.0001, for each condition vs bacterial load). Total conjunctival leucocyte increases (CD45^+^ cells immune infiltrate) also correlated with increases in clinical signs and bacterial loads (p<0.0001, vs both clinical signs and bacterial load). B cell increases correlated with bacterial load changes (p=0.048), and both tear and serum IgG increases (p=0.004 and p=0.0003, for tear and serum respectively). Serum IgA and IgG increases also correlated with increased bacterial loads, while in the tears, only IgG increases were significantly correlated to bacterial load (p=0.034), tear IgA positive correlation was not significant (*p=0.1685*). At the cytokine level, tear MIP1β, MCP-1, IL-8, TGFα, and IL-1RA positively correlated with bacterial load (p<0.0001, p=0.0011, p<0.0001, p=0.0001, and p=0.0002, respectively). In contrast, tear cytokines GM-CSF, to sCD40L (GM-CSF, IL-15, VEGF, IL-5, IL-4, IL-10, IL-1β, IL-12/23, IL-17α, IFN-γ, IL-13, and sCD40L) all correlated with each other, but were all negatively correlated with bacterial load and clinical signs. Altogether, this combined analysis identified a biomarker signature that was indicative of Ct bacterial load and Ct-associated clinical signs, while a separate signature was associated with reductions in bacterial loads and clinical signs.

**Fig 7.**
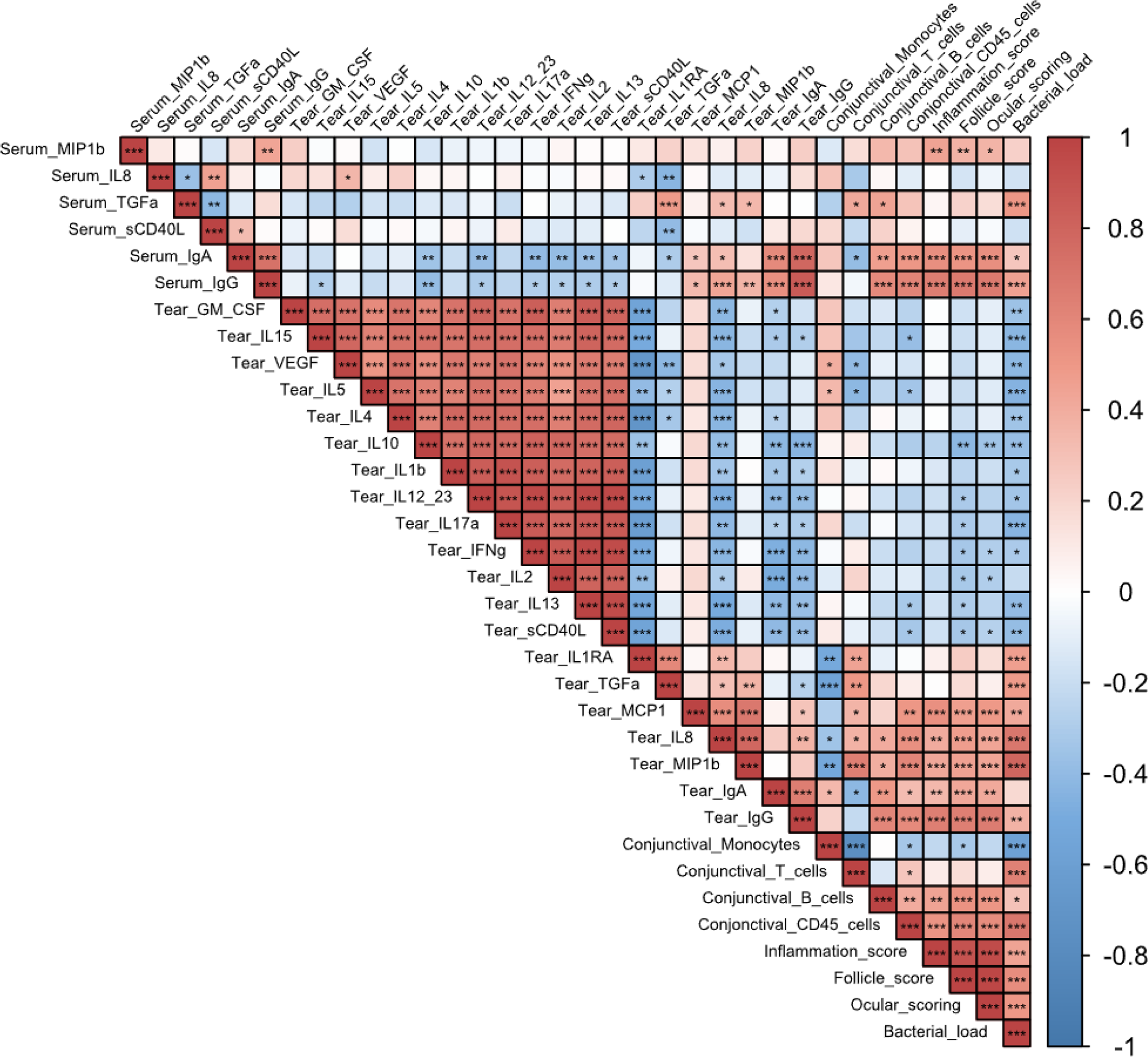
Correlation matrix of all the different parameters measured in group 2 for all time-points. Correlation was done using a non-parametric Spearman correlation test (performed with R software) from -1, maximum negative correlation, to 1, maximum positive correlation. *** p<0.001, ** 0.001<p<0.01, * 0.01<p<0.05.

## Discussion

### A faithful NHP model of trachoma mimicking clinical features of human disease

Similar to previously reported trachoma NHP models [3,12,13], our model faithfully reproduces the clinical features of acute trachoma observed in human patients in endemic populations. In humans, infection of the conjunctiva by Ct is followed by follicular conjunctivitis and conjunctival inflammation, which develop 1-2 weeks p.e. [5]. In our study, Ct inoculation elicited clinical signs in all animals within the same time frame (**Fig 3B**), and their increasing severity was accompanied by increases in conjunctival bacterial loads [4]. In humans, Chlamydial infections are typically cleared within 3 to 8 weeks, but clinical signs of inflammation can persist for 6 to 18 weeks [28], which aligns with experimental findings in our model, since persistence of follicles was observed 11 weeks after bacterial exposure, i.e. 2 to 5 weeks after bacterial clearance.

Given the ongoing burden of trachoma as a public health issue [1,2], it is crucial to have an accurate model for studying the disease [9,11]. Previous studies have used cynomolgus monkeys to reproduce trachoma in NHP. Taylor *et al*. successfully induced conjunctival fibrosis, replicating the full spectrum of the disease [3]. The monkeys were infected on a weekly basis for 52 weeks, highlighting the importance of chronic exposure to Ct for advanced stage disease. Kari *et al*. performed dual Ct exposure, also in cynomolgus monkeys, to test a candidate trachoma vaccine. The second exposure was administered three months after spontaneous clearance of the first inoculation. However, late fibrotic stages of disease were not observed [29]. In our model, a single Ct exposure was employed to characterize clinical and immune makers of acute infection, with the ultimate goal of assessing the efficacy of new vaccine candidates in the future. The novelty of our model is in the development of an optimized clinical grading score (**Fig 2)** [24,30,31], allowing a precise assessment of the kinetics of the clinical disease (**Fig 2**), combined with multimodal exploration of local and systemic immune responses. Optimization of the dose, to observed clinical signs, allowed for precise assessment of clinical disease development kinetics. Clinical signs developed similarly to those seen in humans. Incorporating non-invasive sampling techniques, like conjunctival imprints, proved to be highly valuable in obtaining consistent measurements and establishing novel tools to study kinetics of local immune populations. Furthermore, the analysis of tears encompassed a deeper understanding of the local inflammatory immune landscape in our model. Indeed, our longitudinal *in vivo* data showed similar results to one-time sampling from different trachoma stages in endemic area human cohorts, validating this experimental procedure to faithfully mimic clinical features of human disease [32].

### Two phase immune response: before and after treatment Infectious phase response

Immune infiltrates peaked from weeks 2 to 4 p.e., with a major proportion consisting of T cells. The bacterial load-associated immune response observed in the first 7 weeks was also associated with conjunctival T lymphocytes (*r = 0.63, p < 0.0001*) and T cell survival factors (IL-2 and IL-15) were detected frequently during this period. In humans, T cells seem to play a major role in disease pathogenesis but also are associated with clearance of Ct infection, driven by CD4 T cell-derived IFN-γ [5]. Interestingly in our model, conjunctival T cell numbers did not correlate with T-cell associated cytokines and instead IL-10, IL-2, IL-4, IL-5, IL-17α, and IFN-γ all peaked at week 11 p.e.. The early conjunctival T cell spike was instead associated with cytokines excreted by epithelial cells (IL-1RA, IL-8) and macrophages (MCP-1, MIP1β). In conjunctival swabs from patients, Th1 pro-inflammatory cytokines (IL-1β and TNFα) reveal a strong correlation between acute trachoma and with Ct bacterial load [33]. In contrast, Th1 cytokines evaluated in our panel (IL-1β, TNF-α, and GM-CSF) did not significantly increased p.e. [32].

Additional cytokines that increase in trachoma patients include IL-10 (associated with anti-inflammatory response), CXCL8 (IL-8, involved in recruiting granulocytes and facilitating phagocytosis), and CCL2 (MCP-1, involved in attracting monocytes, lymphocytes, and basophils, as well as promoting macrophage differentiation) [33]. These findings align with our results where IL-8 and MCP-1 significantly increased after Ct exposure.

In animal models, neutrophils have also been shown to have a central role in acute trachoma response. Indeed, Lacy *et al.* showed their involvement in T cell recruitment (both CD4^+^ and CD8^+^) by using neutrophil depletion in a guinea pig model of *Chlamydia cavia* infection. The recruitment of CD4^+^ and CD8^+^ T cells by the influx of neutrophils was a beneficial immune phenomenon that helped to clear the infection [17]. But while this T cell recruitment reduces the bacterial burden, it simultaneously may have an exacerbating effect on ocular symptoms (follicles and inflammation), a tendency also found in a murine model of Ct vaginal infection with concomitant neutrophil depletion [19]. Finally, Lacy *et al*. showed a link between neutrophil depletion and increased IgA titers. This observation could mean that in the absence of neutrophils, while clinical signs are reduced, local IgA increases can occur due to a local increase in B lymphocytes [17]. Similarly, we observed a tendency toward decreased ocular signs around 6 weeks p.e. corresponding to a peak in specific anti-Ct IgA in tears, concomitant with a higher proportion of B cells. Transcriptome studies in humans with active trachoma inferred a prominent role of the innate immune response, notably neutrophils. Natividad *et al.* identified immune infiltrates with immune populations such as T cells, B cells, and a predominance of innate immunity (both NK cells and neutrophils) [34].

In our study, while we did not detect significant neutrophil infiltrates (due to technical limitations), neutrophils in blood increased at the peak of infection (between 2 and 4 weeks p.e.). However, we found a significant T and B cell conjunctival infiltrate (**Fig 4**), that corresponded to follicular lesions, as detected in both humans [20,34] and other animal models [12,17]. In our results, a positive correlation was observed between clinical signs and B lymphocytes infiltrate (r coefficient = 0.52, p = 0.00032). A 1977 study on immunoglobulins in trachoma patients’ tears observed a lower quantity of IgA in trachoma cases than in healthy patients (without significant differences between trachoma phases) [35]. However, Taylor *et al*., in a Cynomolgus model of trachoma, although using another Ct strain (E/Bour), found a delayed increase in tear IgA after conjunctival Ct exposure, and persistence of systemic IgG [23], consistent with our findings.

### Convalescent phase response

The convalescent phase defined as after treatment from 7 weeks p.e., was characterized by an absence of Ct due to either spontaneous clearance (for 4/12 animals) or the effect of azithromycin treatment. The clearance of bacteria coincided with a reduction in conjunctival inflammation, although the presence of follicles persisted, a dissociated pattern that has also been observed in humans [36]. Studies conducted on human cohorts at different stages of trachoma revealed an association between chronic trachoma and the local presence (in mucosal sponges) of IL-1RA (an antagonist of the pro-inflammatory IL-1), as well as IL-4 and IL-13 (associated with Th2 response) [33].

The reduced local leucocytes population at 9 and 11 weeks p.e. could be a result of the higher quantity of IL-1RA at earlier time-points, that may have had an anti-inflammatory effect. In those later time-points, while we observed a reduction in local leucocyte populations, we also saw a higher proportion of monocytes, the cell type mainly responsible for IL-1RA secretion.

In another study involving 470 Tanzanian children, analysis of ocular fluid samples using transcriptome revealed a correlation between acute trachoma and elevated expression of IL-17A. The authors concluded that IL-17A and Th17 responses play a central pro-inflammatory role in trachoma [20]. A transcriptomic study on an Ethiopian cohort of trachomatous patients also suggested a central role of other pro-inflammatory cytokines (including IL-1B, CXCL5 or S100A7) in chronic trachoma [37]. In the present study, the increase in IL-17A was delayed, (week 11 p.e.), along with elevated levels of other pro-inflammatory (VEGF, IL-1B, IL-15, and GM-CSF). Some anti-inflammatory cytokines (IL-10, IL-4, and IL-5) were also elevated but with a delay, suggesting that a balance between both anti- and pro-inflammatory cytokines is needed to achieve disease resolution.

A longitudinal study of a Tanzanian cohort of children identified an association between progressive trachomatous scarring and elevated levels of IL-23, inferred the importance of a Th17 response (through the analysis of IL-23A and PDGFB) [38]. Our observations of the late increase in both IL-12/23 and IL-17A align with those findings. From week 3 p.e., and most drastically weeks 4 and 6, a sharp decrease in total leucocyte counts was observed. During this contraction, the proportion of monocytes increased significantly then subsided p.t.. These observations suggest that local monocytes are characteristic of the continuance of clinical signs of inflammation once bacterial infection resolves. Diminution of the proportion of T cells in the model infection is also consistent with observations in the active stages of trachoma in humans. It has been shown that T cells, particularly CD4^+^ T cells, and the Th2 response correlate with scarring trachoma both locally [33,39] and systemically [40].

### Limitations

The use of conjunctival imprints offers several advantages, such as being non-invasive and allowing for repeated sampling over a short period. However, this technique for harvesting cells can affect their viability. Certain sensitive cells, like neutrophils, may lose their viability during the elution step of the protocol [41]. Therefore, the lack of observed neutrophils by conjunctival imprint cytometry is likely attributed to the technical process rather than their absence. Additionally, imprints will likely capture only the cells present on the surface of the conjunctiva, or those exposed to this sampling site in the context of a lesion; those in dermal conjunctival layers are likely not picked up by this method. Sampling of the conjunctival surface may induce a bias in the type of cells sampled by imprints (bias that would be in favor of immune component of follicles and desquamating cells).

The non-identification of a significant proportion of leucocytes (labeled as “other leucocytes”) could be due to missing markers in our panel (such as markers for dendritic cells or granulocytes populations), technical difficulties (dead cells, non-leucocytes expressing CD45 antigen), or could imply non-specific binding of antibodies [42].

Our current model has laid the groundwork for the characterization of the local immune response, however in order to understand the specific adaptive immune response to ocular infection a deeper analysis of specific T cell response is required.

To summarize, our NHP model of acute trachoma adequately reproduces hallmarks of acute human disease. Most of the changes in immune effectors were observed at the local level, with an influx of T lymphocytes at the peak of infection and the persistence of monocytes with clinical signs in the absence of infection. This acute model could be further developed to investigate chronic trachoma model by repeated Ct inoculations to induce conjunctival fibrosis, for pre-clinical assessment of therapeutic strategies and characterization of immune responses. This model infection may also be of importance in studies of treatment immune-modulation and in the identification of non-invasive correlates of protection for use in human vaccine trials.

## Acknowledgment

This study was supported by the European Commission through the TracVac consortium contract (Grant agreement ID: 733373) and The Innovation Fund Denmark (069-2011-1). The Infectious Disease Models and Innovative Therapies research infrastructure (IDMIT) is supported by the Domaine d’Intérêt Majeur (DIM) “One Health” program, and the “Programme Investissements d’Avenir” (PIA), managed by the ANR under references ANR-11-INBS-0008 and ANR-10-EQPX-02-01. We thank the Thea foundation and Didier Renault for their financial support. We also thank Quentin Sconosciuti, Victor Magneron, Maxime Potier, and Jean-Marie Robert for the NHP experiments; Laetitia Bossevot, Laura Junges, Laurine Moenne-Loccoz and Loïc Pintore for the RT-qPCR assays; Mario Gomez-Pacheco and Jérôme van Wassenhove for the flow cytometry helps; Yann Gorin for the logistics and safety management; Isabelle Mangeot and Camille Morice for help with resources management; Brice Targat and Karl-Stefan Baczowski who contributed to data management and Oscar Haigh for help during manuscript proofreading.

## Authors contributions

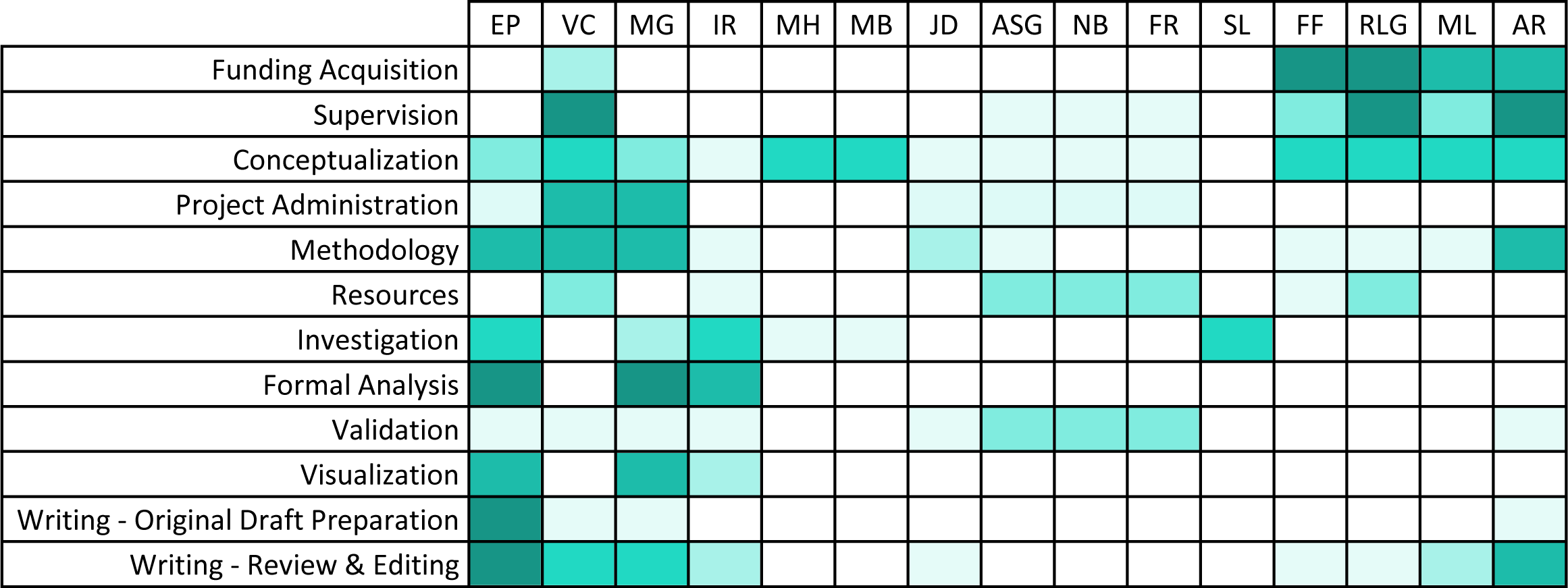

## Supplementary data

**Fig S1.**
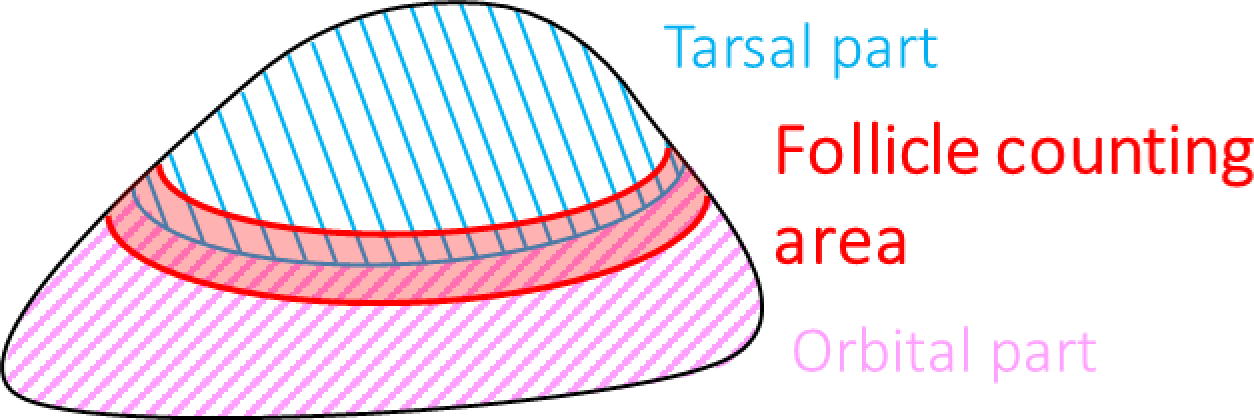
Follicules counting area in the superior tarsal conjunctiva. Follicle counting area is shown in red at the interface of the tarsal part (blue) and the orbital part (pink) of the palpebral conjunctiva.

**Fig S2.**
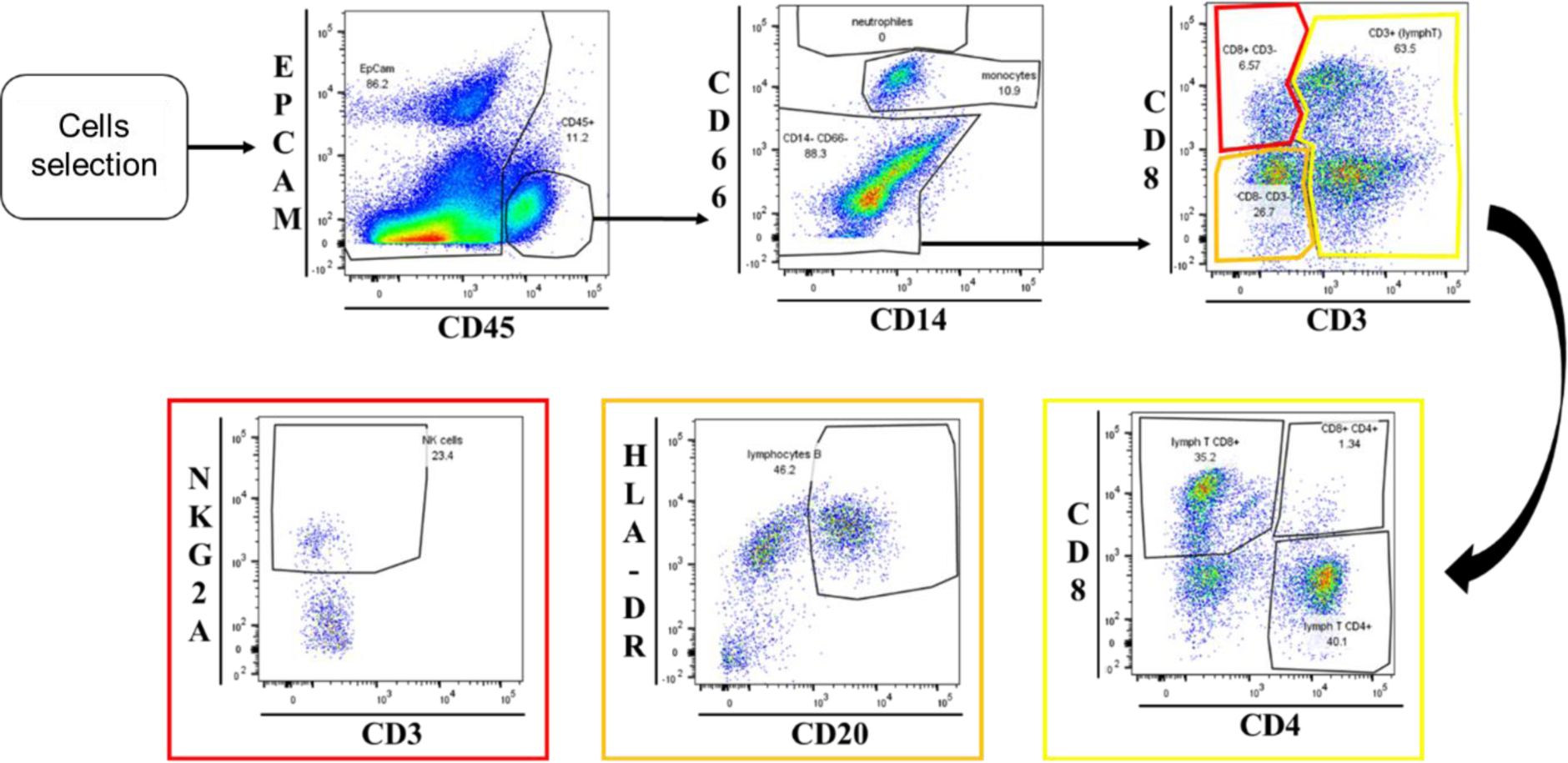
Gating strategy of pseudo-color plots windows performed using FlowJo software and applied to cells eluted from conjunctival imprints. This strategy was used to assess the following ocular surface cell immune populations: leukocytes (CD45^+^), neutrophils (CD66^+^), monocytes (CD14^+^), T cells (CD3^+^), CD4 T cells (CD3^+^, then CD4^+^/CD8^-^), CD8 T cells (CD3^+^, then CD4^-^/CD8^+^), natural killer cells (CD3^-^/CD8^+^, then NKG2A^+^), B cells (CD20^+^ and HLA-DR^+^). This strategy is applied to conjunctival imprints’ cells’ flow cytometry data.

**Fig S3.**
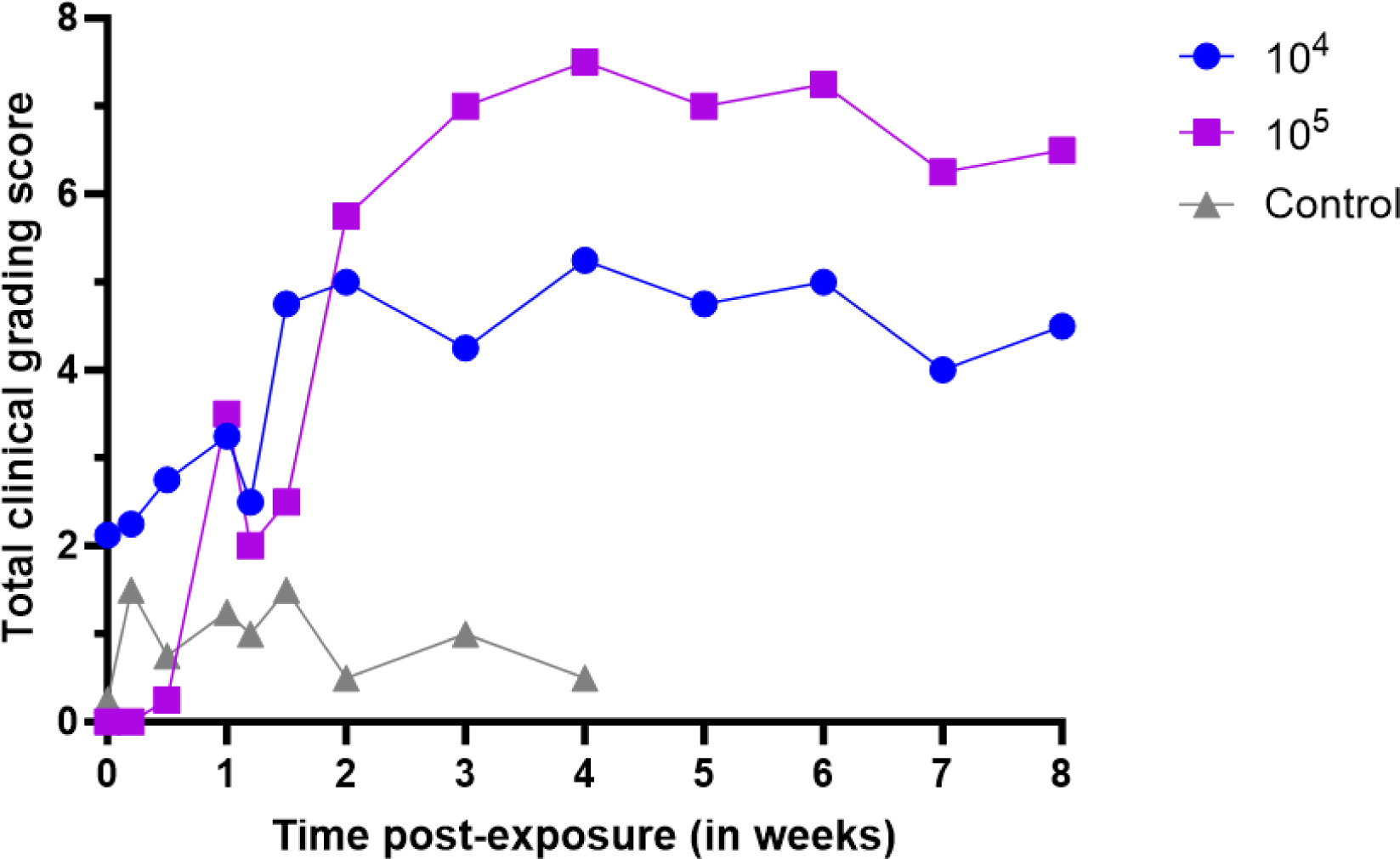
Total Conjunctival clinical scores of Group 1.

**Fig S4.**
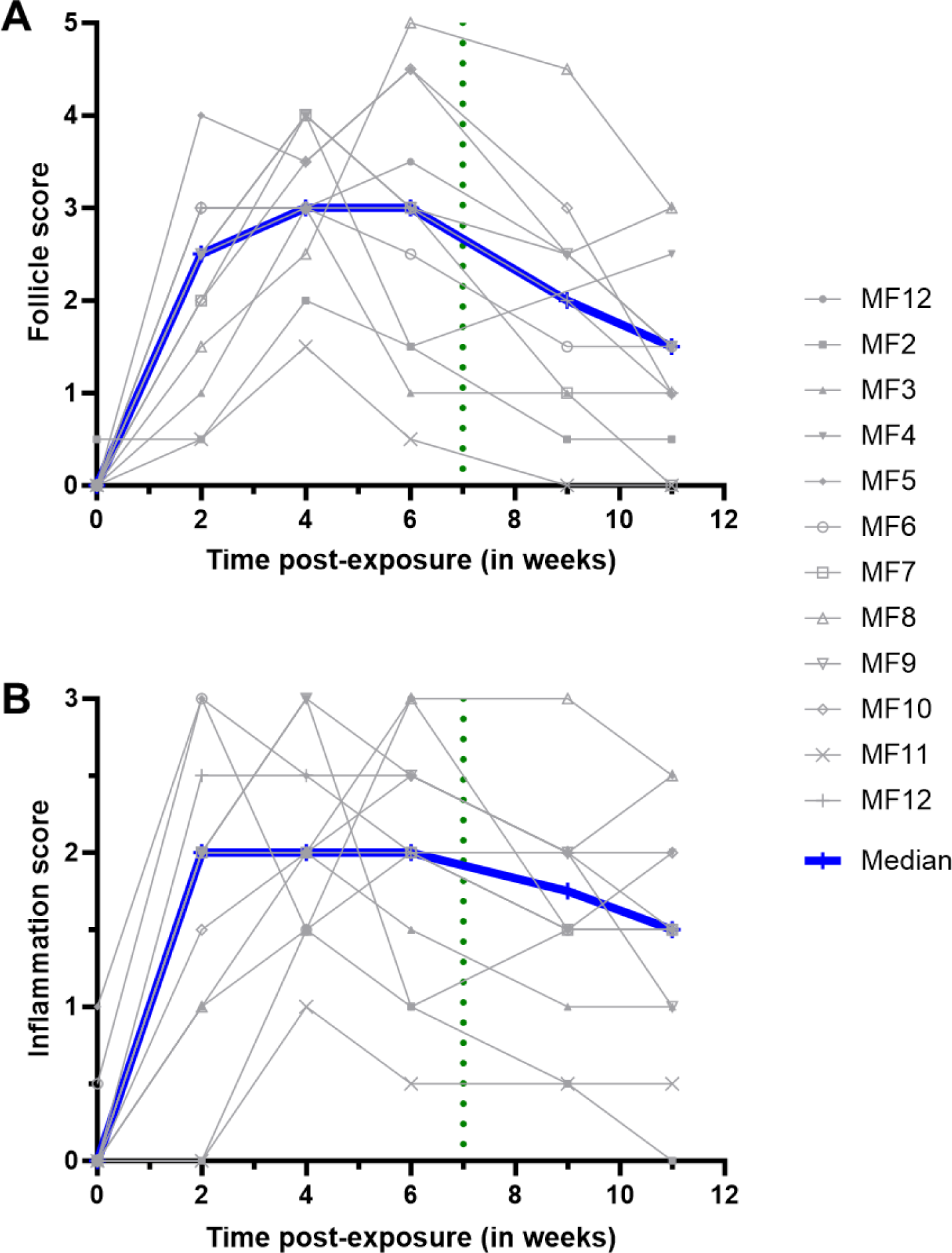
Conjunctival clinical scoring results for follicle score (A) and inflammation (B) in Group 2.

**Fig S5.**
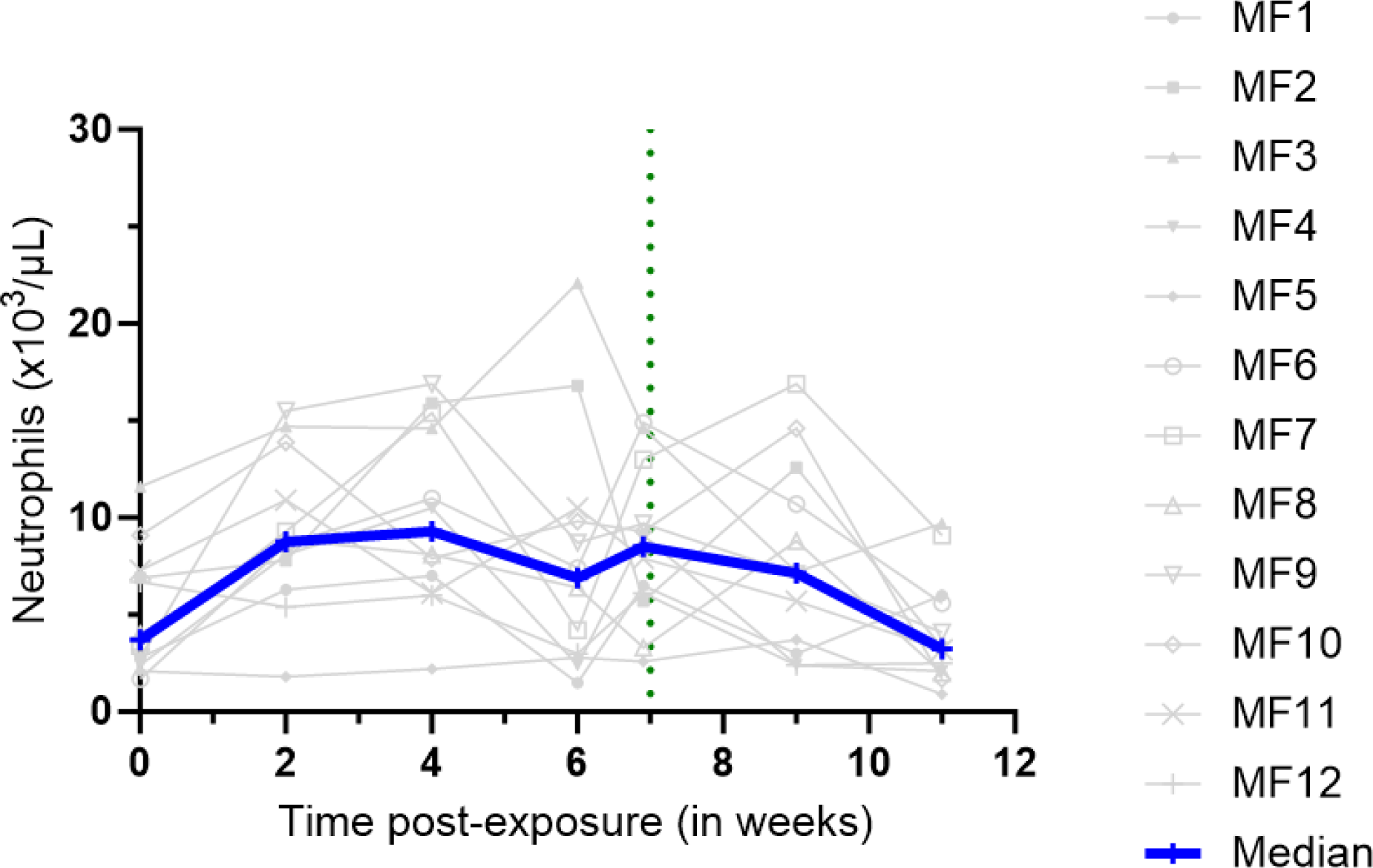
Blood neutrophil count in group 2. (performed at each timepoint as part of a complete blood count.

**Fig S6.**
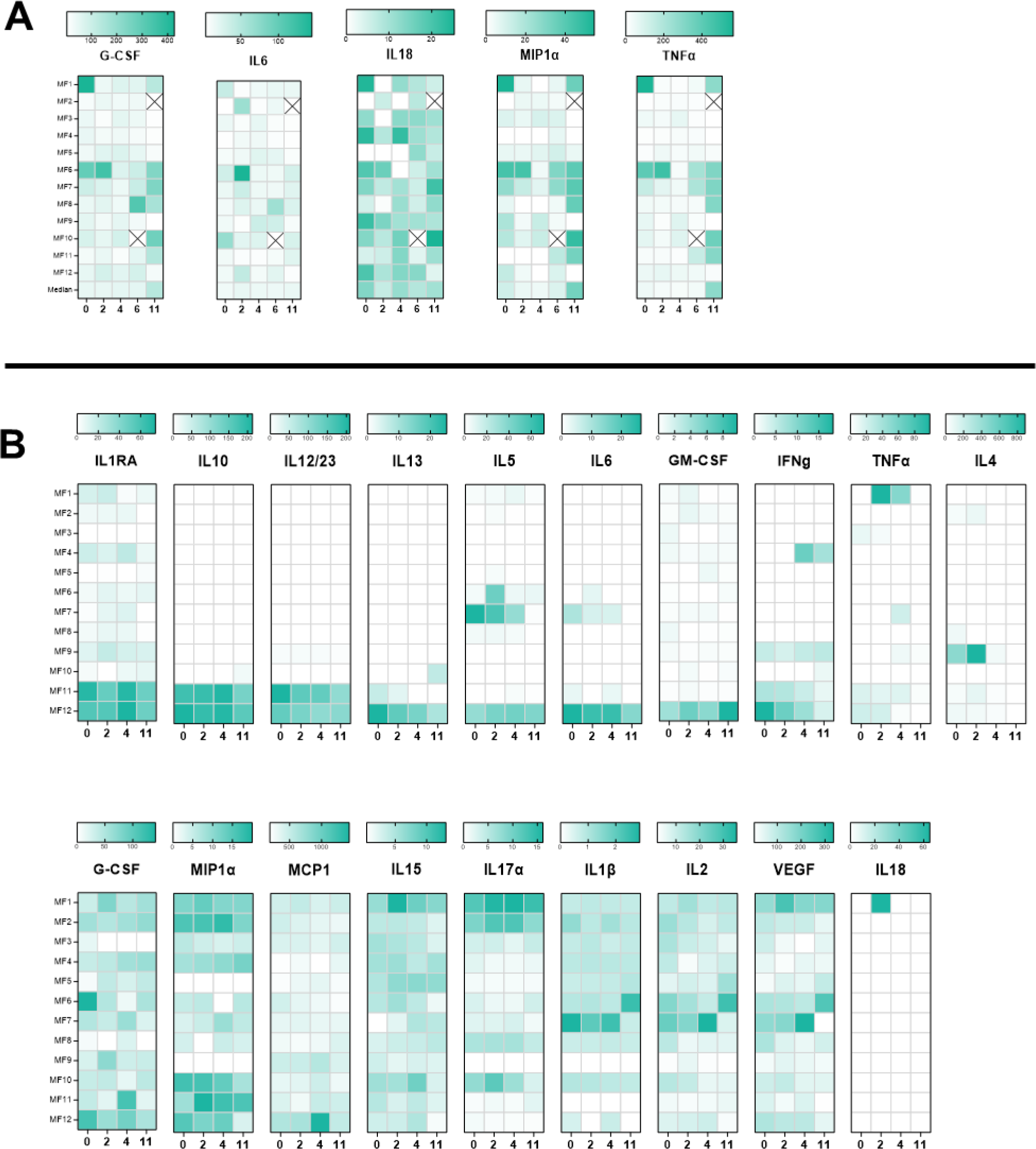
Cytokines quantification without statistically significant changes quantified on tears (A) and serum (B).

## Notes

### Competing Interest Statement

Funding for this work was recieved from the Thea foundation (EP, ML, AR). Additionnally roles of consultants and occasional speakers for the Thea foundation is disclosed (ML, AR). These competing interests do not alter adherence to PLOS policies on sharing data and materials.

